# Joint degeneration in a mouse model of pseudoachondroplasia: ER stress, inflammation and autophagy blockage

**DOI:** 10.1101/2021.06.04.447121

**Authors:** Jacqueline T. Hecht, Alka C. Veerisetty, Mohammad G. Hossain, Debabrata Patra, Frankie Chiu, Francoise Coustry, Karen L. Posey

**Affiliations:** Department of Pediatrics, McGovern Medical School, The University of Texas Health Science Center at Houston (UTHealth), Houston, 77030 TX, USA; School of Dentistry, The University of Texas Health Science Center at Houston (UTHealth), Houston, 77030 TX, USA; Institute of Clinical and Translational Sciences Washington University at St. Louis, St. Louis, 63130 MO, USA

**Keywords:** Cartilage oligomeric matrix protein, autophagy, ER stress, dwarfism, chondrocyte, articular cartilage, joint degeneration

## Abstract

Pseudoachondroplasia (PSACH), a short limb skeletal dysplasia, associated with premature joint degeneration is caused by misfolding mutations in cartilage oligomeric matrix protein (COMP). Here, we define mutant-COMP-induced stress mechanisms that occur in articular chondrocytes of MT-COMP mice, a murine model of PSACH. The accumulation of mutant-COMP in the ER occurred early in MT-COMP articular chondrocytes and stimulated inflammation (TNFα) at 4 wks. Articular chondrocyte death increased at 8 wks and ER stress through CHOP was elevated by 12 wks. Importantly, blockage of autophagy (pS6), the major mechanism which clears the ER, sustained cellular stress in MT-COMP articular chondrocytes. Degeneration of MT-COMP articular cartilage was similar to that observed in PSACH and was associated with increased MMPs, degradative enzymes. Moreover, chronic cellular stresses stimulated senescence. Senescence-associated secretory phenotype (SASP) may play a role in generating and propagating a pro-degradative environment in the MT-COMP murine joint. The loss of CHOP or resveratrol treatment from birth preserved joint health in MT-COMP mice. Taken together, these results indicate that ER stress/CHOP signaling and autophagy blockage are central to mutant-COMP joint degeneration and MT-COMP mice joint health can be preserved by decreasing articular chondrocyte stress. Future joint sparing therapeutics for PSACH may include resveratrol.

## Introduction

Cartilage oligomeric matrix protein (COMP) is a large, matricellular protein that mediates a variety of cell-cell and cell-matrix interactions [1–7]. COMP interacts with many extracellular matrix (ECM) proteins including, but not limited to, collagens types I, II, IX, XII, XIV, matrilin-3, aggrecan and fibronectin [8–12] and may provide a hub for interaction(s) of collagens with proteoglycans and other ECM proteins [8, 9, 11]. COMP likely plays a role in mechanical strength of ECM tissues, as loading increases COMP levels in tendons and aging or overuse decreases abundance [11, 13]. Chondrocyte proliferation and chondrogenesis is stimulated by COMP [14, 15]. Mutations in COMP cause pseudoachondroplasia (PSACH), a severe dwarfing condition characterized by disproportionate short stature, short limbs, joint laxity, pain and early onset joint degeneration [2, 9, 16–38]. PSACH birth parameters are normal and the first sign of the disorder is decelerating linear growth, starting by the end of the first year, and a waddling gait developing by two years of age [29, 32]. Radiographic examination leads to a diagnosis and the key characteristics are shortening of all the long bones, small abnormal epiphyses, widened and irregular metaphyses, small, underossified capital femoral epiphyses and platyspondyly [21, 29, 32, 36, 37]. While loss of linear growth is the most obvious untoward outcome in PSACH, joint dysfunction and pain are the most debilitating and long-term complications. Pain is significant and begins in childhood, likely from the inflammation which plays a role in the underlying growth plate chondrocyte pathology [17, 21, 34, 39, 40]; whereas, the pain in adulthood may reflect joint degenerative changes that necessitate hip replacement in a majority of adults [29]. Joint degeneration in PSACH occurs very early in life between the second or third decades and all joints are affected especially the hips, elbows, and shoulders [29, 37, 41, 42].

Long before COMP mutations were identified as the cause of PSACH, it was known that growth plate chondrocytes contained massive amounts of material in the rER cisternae [43]. Later studies proved that mutant-COMP does not fold properly and, therefore, is retained in the ER [19, 44]. Moreover, mutant-COMP prematurely interacts with binding partners in the ER forming and an ordered matrix composed of types II and IX collagen and matrilin 3 (MATN3) and other ECM proteins, resulting in intracellular protein accumulation [31, 39, 45]. This material is not degraded efficiently enough to maintain chondrocyte function and fills the cytoplasmic space becoming toxic to growth plate chondrocytes [2, 21, 33, 35]. To study the cellular mechanism involved in the PSACH pathology, a doxycycline (DOX)-inducible mouse that expresses mutant-D469del-COMP (designated the MT-COMP mouse) in chondrocytes was generated [39]. The MT-COMP mouse mimics the clinical phenotype and chondrocyte PSACH pathology [39, 46, 47]. Studies of the MT-COMP mouse showed that mutant-COMP is retained in the rER of growth plate chondrocytes stimulating ER stress through the CHOP pathway that in turn activates oxidative and inflammatory processes initiating a self-perpetuating stress loop involving oxidative stress and inflammation. This stress loop leads to DNA damage, necroptosis and loss of growth plate chondrocytes [46–48]. ER stress and TNFα-driven inflammation increase mTORC1 signaling [49] that represses autophagy eliminating a critical mechanism needed to clear misfolded proteins and results in intracellular accumulation in growth plate chondrocytes [49]. This molecular mechanistic process explains the deceleration in linear growth associated with PSACH.

In this work, we focused on delineating the effect of mutant-COMP on joint health in our PSACH mouse model and the timing of joint degeneration. Articular and growth plate cartilages serve very different functions and these tissues mature at different times [50, 51]. Growth plate chondrocytes generate a large volume of matrix driving long-bone growth and after reaching hypertrophy chondrocytes die. In contrast, articular chondrocytes synthesize matrix to provide lubrication and a cushion to withstand compressive forces during movement, and are very long lived. Growth is essential to both fetal and postnatal periods, while withstanding weight bearing compressive forces during ambulation occur postnatally. Both articular and growth plate chondrocytes synthesize different extracellular matrix proteins unique to each tissue. In this study, articular cartilage was evaluated for ER stress, inflammation, autophagy block, proteoglycans and pro-degradative processes and the timing of each process. While stresses that drive mutant-COMP pathology in growth plate chondrocytes also underly the articular cartilage pathology, important unique degradative processes were present.

## Results

### Mutant-COMP is retained in the ER in adult articular chondrocytes

Intracellular retention of mutant-COMP in chondrocytes is a hallmark of PSACH [21, 41, 52]. Previously, we have shown that mutant-COMP accumulates in the ER of growth plate chondrocytes of MT-COMP mice beginning at E15 [39]. **Figure 1** left panel, shows that human mutant-COMP (red signal) is retained in the ER (green signal) of MT-COMP articular cartilage of MT-COMP chondrocytes (yellow merge) at 4, 8, 12, 16, and 20 wks (**F, J, N, R, V**). In contrast, there is no detectable mutant-COMP in control C57BL\6 mice (**Fig. 1A, E, I, M, Q, U**).

**Figure 1:**
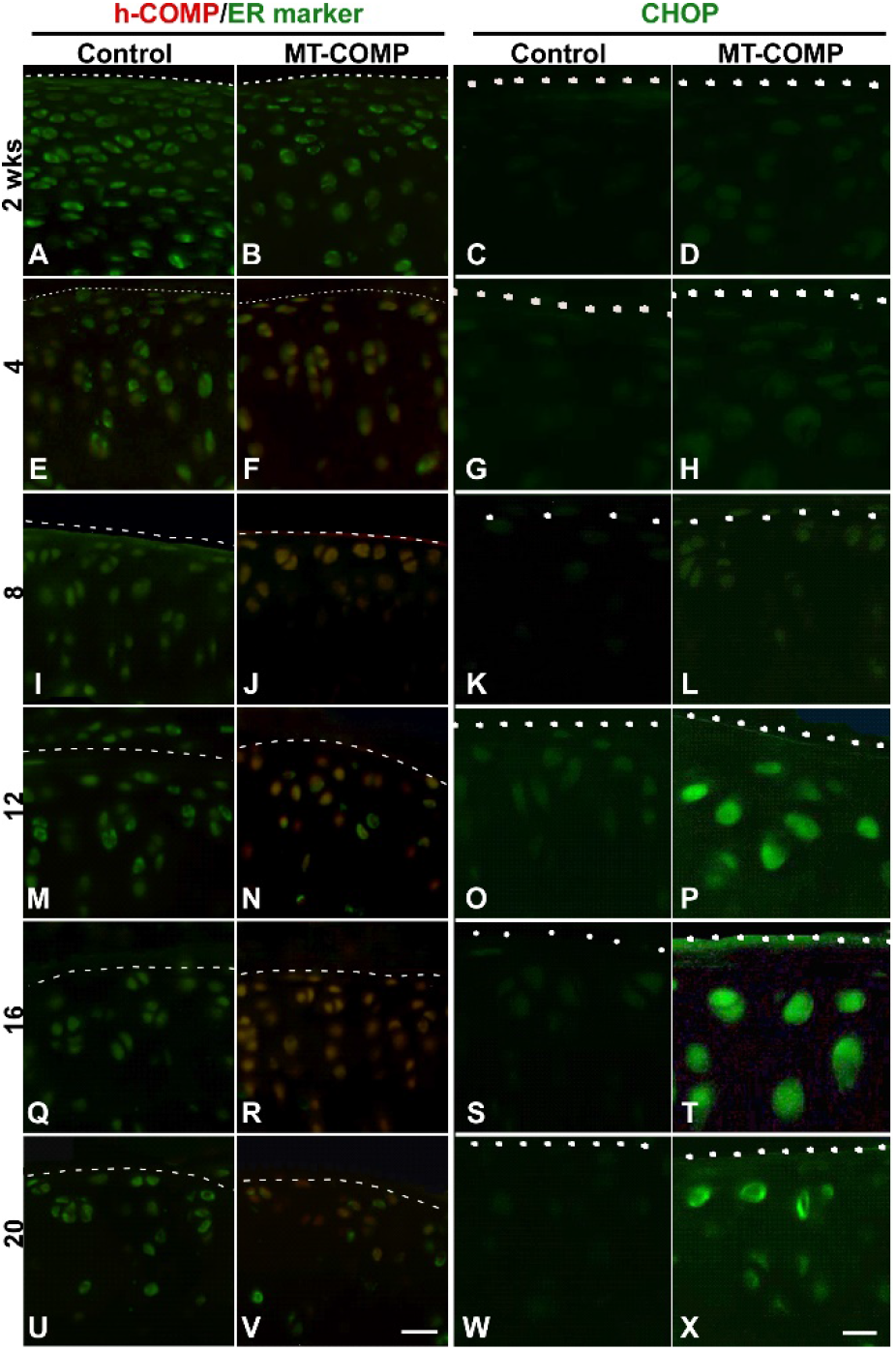
Retention of human mutant-COMP in ER of MT-COMP articular chondrocytes at 4 wks with CHOP expression at 12 wks. DOX administered from birth to limb collection at 2, 4, 8, 12, 16, 20 wks. Control and MT-COMP tibial articular cartilages were immunostained using human COMP-specific antibodies (red) and protein disulfide isomerase (PDI) (green) an ER marker (**A-B, E-F, I-J, M-N, Q-R, U-V**). Mutant-COMP was expressed and retained in ER of articular chondrocytes of MT-COMP at 4 wks (**F, J, N, R, V,** Bar = 100 μm) but not in controls (**A**, **E, I, M, Q, U**). CHOP immunostaining is shown in **C-D, G-H, K-L, O-P, S-T, W-X**. CHOP was present in MT-COMP articular chondrocytes at 12, 16 and 20 wks (**P, T, X)** and absent in controls (**C, G, K, O, S, W**). **A-B, E-F, I-J, M-N, Q-R, U-V** Bar = 100 μm; **C-D, G-H, K-L, O-P, S-T, W-X** Bar = 50 μm.

### ER stress, inflammation, matrix degradation, autophagy repression, senescence and chondrocyte death are present in MT-COMP articular chondrocytes

In growth plate chondrocytes, the intracellular accumulation of mutant-COMP stimulates a complex pathological process. This involves activation of the ER stress through the CHOP pathway, which in turn activates oxidative and inflammatory processes that exacerbates ER stress, causing an over activation of mTORC1 signaling blocking autophagy, and ultimately chondrocyte death [46, 47]. As shown in **Figure 1**, CHOP expression in the MT-COMP articular cartilage is observed at 12, 16, 20 wks of age (CHOP panel-**P, T, X**), while absent in controls (**C,G,K,O,P,S,W**). CHOP is a pro-apoptotic transcription factor that stimulates cell death when ER stress is unresolved [53]. The presence of CHOP in articular chondrocytes at 12 wks demonstrates that ER stress is delayed by 8 wks after the intracellular accumulation of mutant-COMP that begins at 4 wks.

Since inflammatory processes involving pro-inflammatory cytokines, IL-1β and TNFα, play a significant role in the growth plate, these cytokines were investigated in the articular cartilage (IL-1β data not shown). Shown in **Figure 2**, TNFα was evident in the deep zone of articular cartilage at 2 wks of age and was observed in all zones from 4-20 wks compared to little or no signal in controls. TNFα and IL-1β promote the synthesis of matrix metalloproteinases (MMPs), a family of degradative enzymes that cleave collagens and proteoglycans in the extracellular matrix [54]. MMP13 is synthesized by articular chondrocytes and plays an important role in ECM degradation of articular cartilage associated with osteoarthritis (OA) [54–56]. MMP13 immunostaining was increased in MT-COMP mice with minimal staining at 8 wks and strong signal from 12 - 20 wks, compared to control articular cartilage that showed minimal immunostaining (**Fig. 2**). Consistent with these findings MMP (−2, −3, −9, and −13) activity measured at 12 wks by MMPSense was significantly higher in MT-COMP knees (4.26 X 10^8^ ± 6.52 X 10^7^) than controls (3.28 X 10^8^ ± 1.88 X 10^7^) (P< 0.034911) (data not shown). These findings show that joint degeneration-associated cytokines (TNFα beginning at 4 wks) and elevated MMP13 immunostaining were associated with mutant-COMP accumulation in articular chondrocytes in MT-COMP mice.

**Figure 2:**
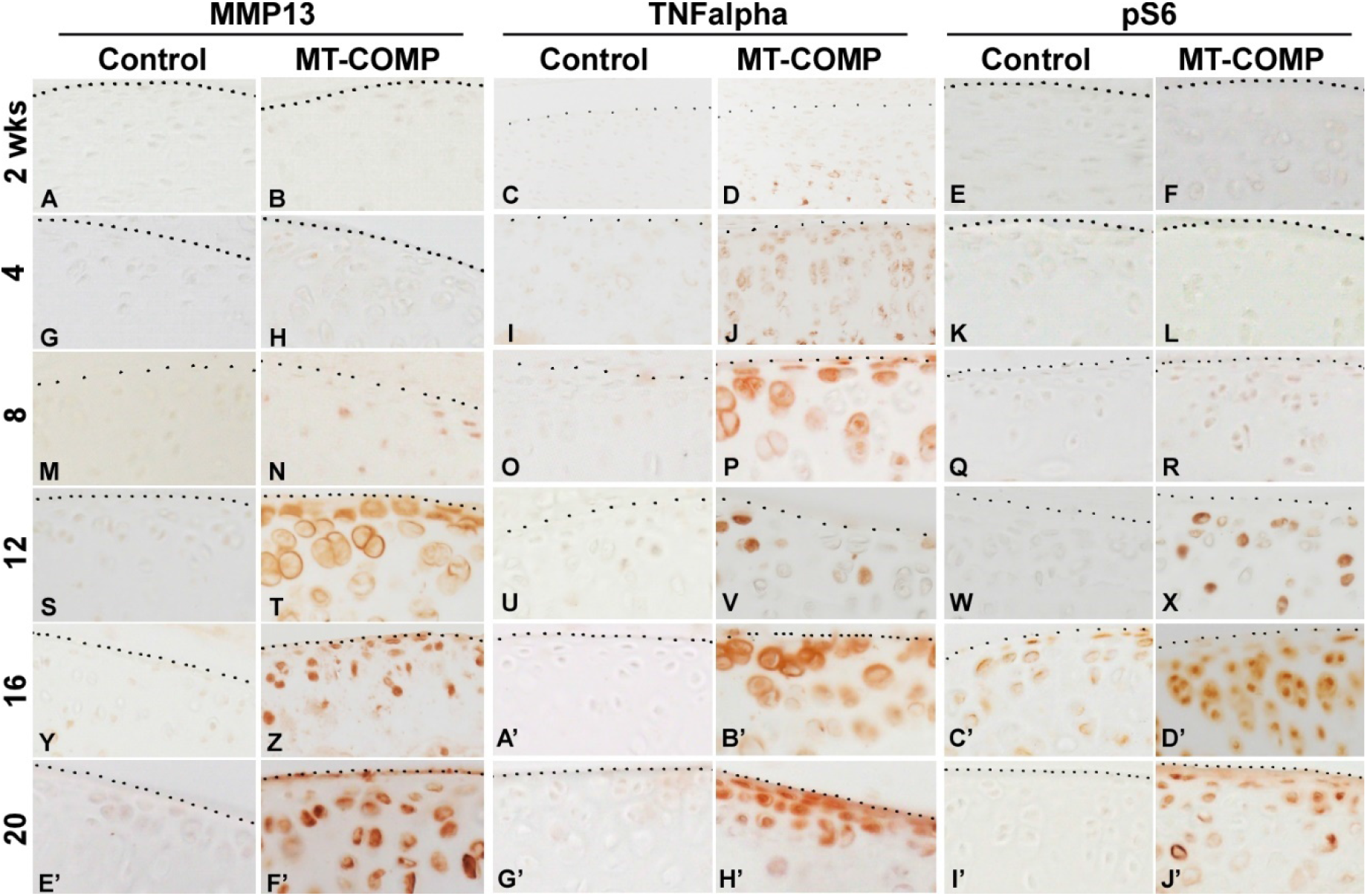
Degradation, inflammation and autophagy blockage is increased in MT-COMP articular cartilage/chondrocytes. DOX was administered from birth to collection at 2, 4, 8, 12, 16, 20 wks. Control and MT-COMP tibial articular cartilage were immunostained using MMP13 (**A-B, G-H, M-N, S-T, Y-Z, E’-F’**), TNFα (**C-D, I-J, O-P, U-V, A’-B’, G’-H’**) or pS6 (**E-F, K-L, Q-R, W-X, C’-D’, I’-J’**). Control mice show no MMP-13 or TNFα signal and minimal pS6. In contrast, the MT-COMP articular chondrocytes show minimal MMP-13 at 8 wks that is increased by 12 wks; TNFα expression is found in the deep zone of the articular cartilage at 2 wks of age and in all three layers at 4, 8, 12, 16, 20 wks; pS6 signaling is seen at 12, 16, and 20 wks. Bar =

Autophagy is repressed in both, OA articular chondrocytes [57] and MT-COMP growth plate chondrocytes [49], therefore, MT-COMP articular chondrocytes were assessed for the presence of phosphorylated S6 ribosomal protein (pS6). pS6 is an established readout for mTORC1 signaling which regulates synthesis of cellular components and inhibits autophagy [58]. As shown in **Figure 2**, pS6 signal was increased in MT-COMP articular chondrocytes beginning at 12 wks, indicating autophagy suppression. In addition to autophagy blockage, articular chondrocyte death increased beginning at 8 wks (**Fig. 3**) and the loss of these chondrocytes may impact matrix synthesis. Moreover, the increase in CHOP expression proceeds the peak in articular chondrocyte cell death at 16 wks suggesting that CHOP contributes to chondrocyte death in MT-COMP mice (**Fig. 1** CHOP panel and **Fig. 3**).

**Figure 3:**
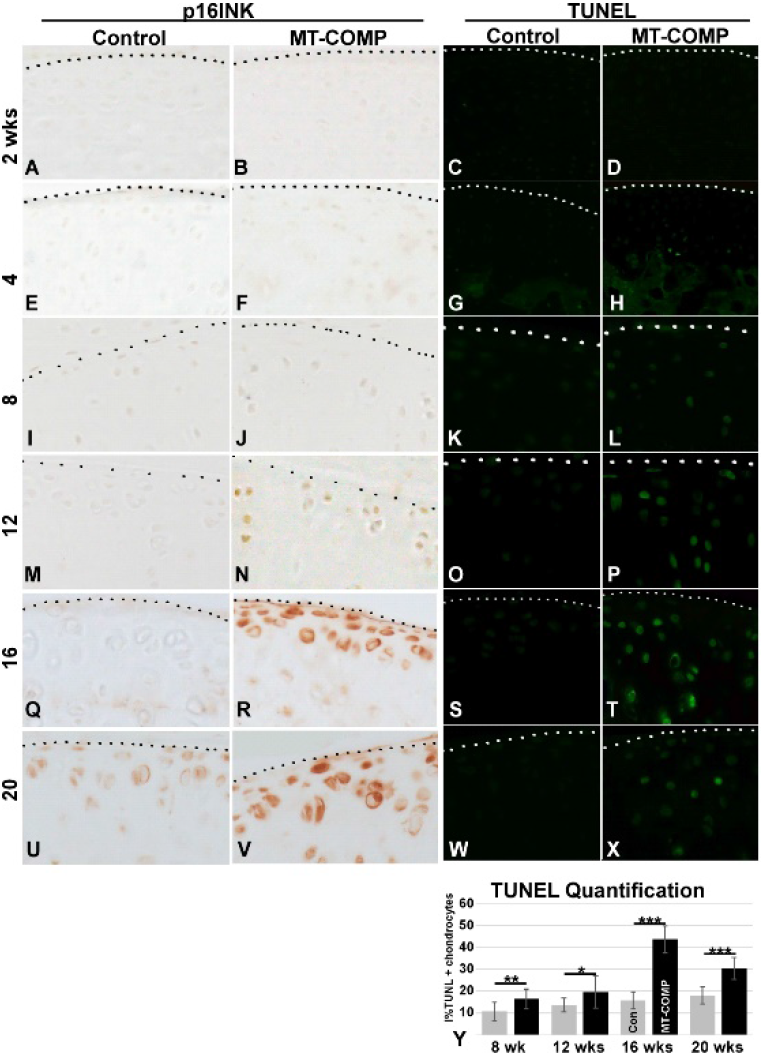
Articular chondrocyte senescence and death in MT-COMP mice. DOX administered from birth to collection at 2, 4, 8, 12, 16, 20 wks. Control and MT-COMP tibial articular cartilage were immunostained using p16 INK4s antibodies (**A-B, E-F, I-J, M-N, Q-R, U-V**). p16 INK4a expression is present from 12-20 wks (**N, R, V**). The MT-COMP mice show numerous TUNEL positive articular chondrocytes (**L, P, T, X**) compared to controls (**C, G, K, O, S, W)**. Percent TUNEL positive MT-COMP articular chondrocytes shown at 8, 12, 16, 20 wks. MT-COMP TUNEL staining is significantly different than controls*P≥0.05; **P≥0.005; ***P≥0.0005 (**Y**). Sections from the hind limb articular cartilage of at least 10 animals per group were stained with TUNEL. Bar =

Senescence is known to play a role in OA joint degeneration [59, 60] and was evaluated using senescence marker p16 INK4a. As shown in **Figure 3**, p16 INK4a was observed in MT-COMP articular chondrocytes from 16-20 wks, consistent with the timing of joint degeneration in MT-COMP mice (**Fig. 4**) [59, 60]. This suggests that MT-COMP articular chondrocytes experience cellular stress which corresponds with senescent articular chondrocytes by 16 wks, a finding typically associated with aging and not with relatively young adult mice [61–63].

**Figure 4:**
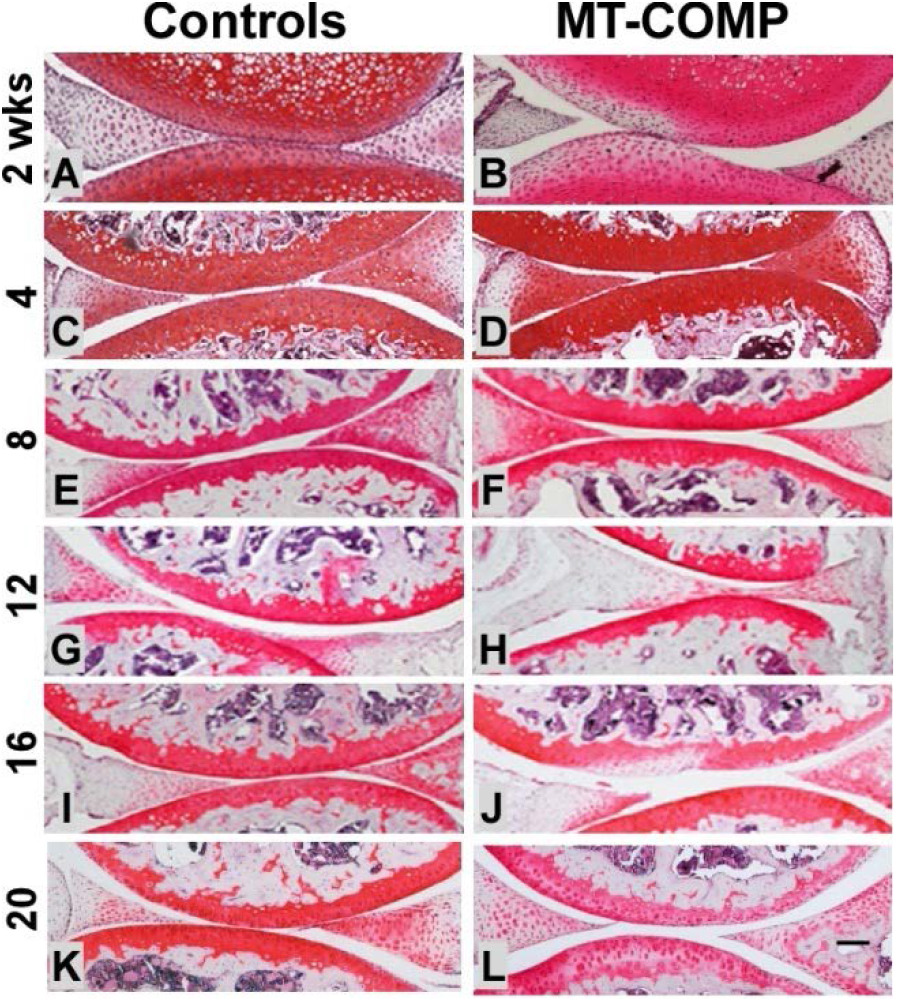
Reduced proteoglycans in MT-COMP mice by 16 wks. DOX administered from birth to collection at 2, 4, 8, 12, 16, 20 wks. Safranin O staining of control and MT-COMP at 2, 4, 8, 12, 16 and 20 wks. Abundant and rich proteoglycan layer was found in control articular cartilage (**A, C, E, G, I, K**). The proteoglycan layer in MT-COMP mice was similar to controls at 4 and 8 wks (**D, F**) but was diminished in select areas at 16 wks (**J**) and progressed to a more generalized loss at 20 wks (**L**). Bar = 500 μm.

### Decreased proteoglycans in MT-COMP articular cartilage at 16 wks

MT-COMP and control joints were stained with Safranin O at 2, 4, 8, 12, 16 and 20 wks to assess proteoglycan content because proteoglycan loss is an early marker of joint degeneration [64–68]. As shown in **Figure 4,** control articular cartilage has a dense layer of proteoglycans at all ages (**A, C, E, G, I, K**). In contrast, MT-COMP articular cartilage shows depletion of proteoglycans beginning at 16 wks and becoming more pronounced at 20 wks (**Fig. 4J and L**). These findings indicate that the MT-COMP mice have premature joint degeneration similar to that observed in PSACH [32, 37, 42].

### MT-COMP mice show signs of pain in early adulthood

In mice, the presence of pain is measured by changes in specific behaviors [69]. One such behavior is changes in voluntary running. The mouse is housed in a cage with a wheel and the number of rotations are recorded to measure voluntary running based on the assumption is that pain with physical activity will reduce the number of wheel rotations [70]. **Figure 5A** shows that MT-COMP mice ran ≈50% less than controls from 20 to 24 wks of age suggesting that MT-COMP mice are less motivated to run, presumabley due to pain with ambulation.

**Figure 5:**
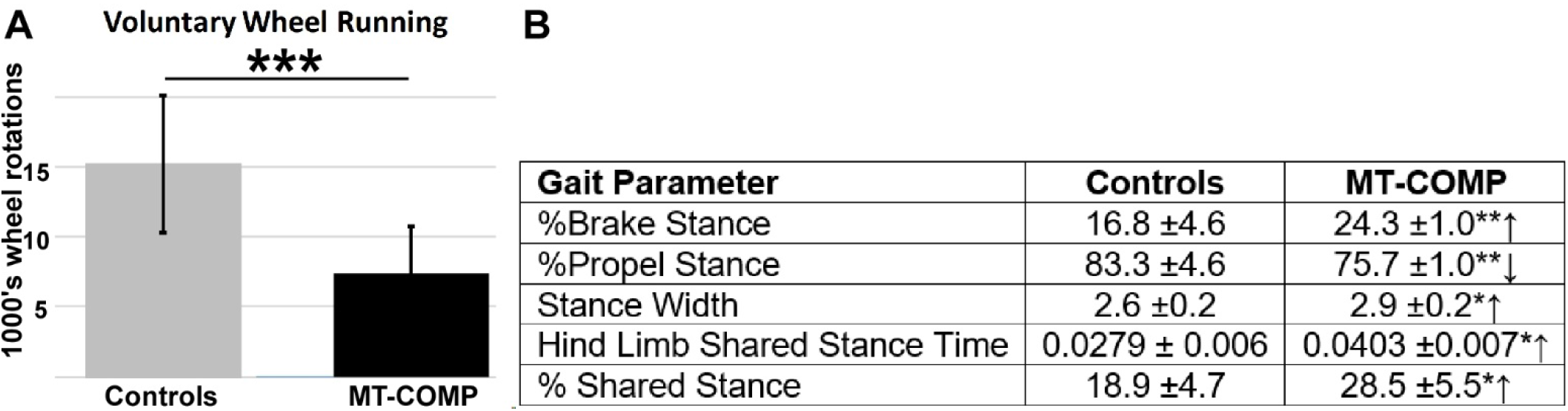
Voluntary running and gait indices. DOX administered from birth to collection. Voluntary running from 20-24 wks (A) and gait indices (B) measured at 16 wks. MT-COMP mice ran approximately 50% less than controls (N= 6/group). Select gait parameters associated with pain are shown and were significantly different in MT-COMP mice compared to controls (N= 6/group). MT-COMP mice had increased %brake stance, absolute paw angle, stance width, hind limb shared stance time and %shared stance. <u>%Brake Stance</u> = the percentage of the stance spent in braking; <u>%Propel Stance</u> = the percentage of the stance spent in propulsion; <u>Absolute Paw Angle</u> = Absolute value of the paw angle; <u>Stance Width</u> = the perpendicular distance between the centroids of either set of axial paws during peak stance; <u>Step Angle</u> = the angle made between left and right hind paws as a function of stride length and stance width; <u>Hind Limb Shared Stance Time</u> = time both hind paws contact belt; <u>%Shared Stance</u> = the percentage of the stance spent with both hind paws contact the belt. *P≤0.05; **P≤0.005; ***P≤0.0005.

Alterations of gait can indicate the presence of pain because these alteration are a natural attempt to reduce stress on limbs associated pain during ambulation [71]. The digigait system is a treadmill that measures multiple gait parameters and can uncover subtle changes in gait. Gait was evaluated at 16 wks by running at 30 cm/sec 15° downhill. As shown in **Figure 5B**, several gait indices that suggest pain were altered including: % brake stance; % propel stance; stance width and hind limb shared stance time. MT-COMP mice spent more time braking and less time in the propel phase of stance. Longer duration in braking stance may indicate more precise control and distribution of load to reduce peak loading [72]. Importantly, hind limb shared stance time and % shared stance time were increased in MT-COMP mice compared to controls (**Fig. 5B**). Hind limb shared stance time is the amount of time that both limbs are in contact with the surface. Shared stance time increases with joint pain because distribution of body weight over both limbs reduces pain [73]. Overall these measures of gait disturbance suggest that MT-COMP mice have a wider gait for stability, care is taken when placing paws on the belt and more time with both hind limbs on the belt to minimize stress on the limbs suggestive of pain.

### Joint degeneration in MT-COMP mice is validated by OA scoring

OA scoring was used to detect and quantify early degenerative changes in the proteoglycan content of articular cartilage, synovitis and bone/cartilage. No differences were detected prior to 16 wks of age. MT-COMP femur proteoglycan content was less and a trend toward higher score for bone/cartilage damage and total score at 16 wks (**Table 1**). Total score of 5.7 was significantly higher in 20 week MT-COMP mice compared to 2.56 in controls (with maximum score of 12). This score included the cartilage/bone damage score in 20 week MT-COMP mice of 0.9 compared to 0.11 in controls and the synovitis score of 1.8 compared to 0.89, respectively (**Table 1**). These findings show that joint degeneration occurs in MT-COMP mice by 20 wks, much earlier than occurs in the C57BL\6 control background strain that occurs after 1 year [74], again consistent with early joint degeneration observed in PSACH.

**Table 1:** OA score for MT-COMP and control joints. Each category is: Pro. Femur = proteoglycan content in femur articular cartilage; Pro. Tibia = proteoglycan content in tibial articular cartilage, Bone/Cartilage and Synovitis is scored from 0-3 with maximum total of 12. Average in each category ± standard deviation is presented with significant p-values *P≤0.05; **P≤0.005 in bold and trending p-values < 0.10 are also shown.

| **Genotype-age wks** | **Pro. Femur** | **Pro. Tibia** | **Bone/Cartilage** | **Synovitis** | **Total** |
| --- | --- | --- | --- | --- | --- |
| Control 4wks | 0.78±0.63 | 0.56±0.50 | 0.11±0.31 | 0.33±0.47 | 1.78±1.13 |
| MT-COMP 4wks | 0.70±0.64 | 0.30±0.46 | 0.40±0.66 | 1.0±0.89 | 2.4±1.85 |
| P value | - | - | - | - | - |
| Control 8wks | 0.80±0.60 | 0.60±0.66 | 0.10±0.30 | 0.40±0.66 | 1.90±1.76 |
| MT-COMP 8wks | 1.3±0.90 | 0.63±0.14 | 0.40±0.49 | 0.60±0.92 | 2.28±0.11 |
| P value | - | - | - | - | - |
| Control 12wks | 0.77±0.32 | 0.57±0.26 | 0.30±0.20 | 0.52±0.30 | 1.48±1.06 |
| MT-COMP 12wks | 1.10±0.83 | 1.20±1.10 | 0.90±0.94 | 1.00±0.89 | 4.20±3.25 |
| P value | - | - | - | - | - |
| Control 16wks | 1.00±0.63 | 0.80±0.60 | 0.50±0.50 | 0.90±0.83 | 3.20±2.27 |
| MT-COMP 16wks | 1.60±0.49 | 1.10±0.54 | 1.10±0.83 | 1.50±0.81 | 5.30±2.19 |
| P value | **0.0372\*** | - | 0.0798 | - | 0.0614 |
| Control 20wks | 1.00±0.78 | 0.56±0.65 | 0.11±0.32 | 0.89±0.83 | 2.56±2.15 |
| MT-COMP 20wks | 1.80±0.87 | 1.20±0.60 | 0.90±0.70 | 1.80±0.60 | 5.70±2.14 |
| P value | 0.0683 | 0.0542 | **0.0091\*\*** | **0.0218\*** | **0.0133\*** |

### Prevention of joint degeneration in MT-COMP mice with resveratrol treatment or ablation of CHOP

Previously, we have shown that resveratrol treatment reduces ER stress by promoting autophagy that clears accumulated mutant-COMP from the ER in growth plate chondrocytes of MT-COMP mice [75]. Resveratrol relieved the ER protein accumulation and reduced chondrocyte death, restoring proliferation and supporting limb growth in juvenile MT-COMP mice [75]. Based on these results, resveratrol was administered to determine whether it could prevent and/or reduce joint degeneration in MT-COMP mice. As shown in **Figure 6,** resveratrol treatment from birth to 20 wks dramatically reduced the loss of proteoglycans in the articular cartilage of adult mice (**Fig. 6A-D**). Importantly, resveratrol treatment prevented the accumulation of mutant-COMP in the ER of articular chondrocytes (**Fig. 6E-H**) reduced ER stress (**Fig. 6M-P**), substantially reduced articular chondrocyte death (**Fig. 6I-L**) and TNFα inflammation (**Fig. 6Q-T**). Consistent with normalization of proteoglycans in the articular cartilage of MT-COMP mice treated with resveratrol, MMP-13 signal was markedly reduced (**Fig. 6Y-B’**). Resveratrol reduced multiple mutant-COMP pathologies including inflammation, ER stress and importantly block of autophagy which allow the chondrocytes to clear misfolded protein and prevent cell death. Similarly, genetic ablation of CHOP was used to interrupt the ER stress signaling pathway that breaks down the pathological loop between ER stress, inflammation and oxidative stress. MT-COMP/CHOP^-/-^ articular cartilage at 20 wks was healthy (**Fig. 6D**) and MMP-13 was minimal (**Fig. 6B’**). The loss of CHOP considerably reduced accumulation of mutant-COMP in articular chondrocytes (**Fig. 6H**) and the number of TUNEL positive chondrocytes (**Fig. 6L**).

**Figure 6:**
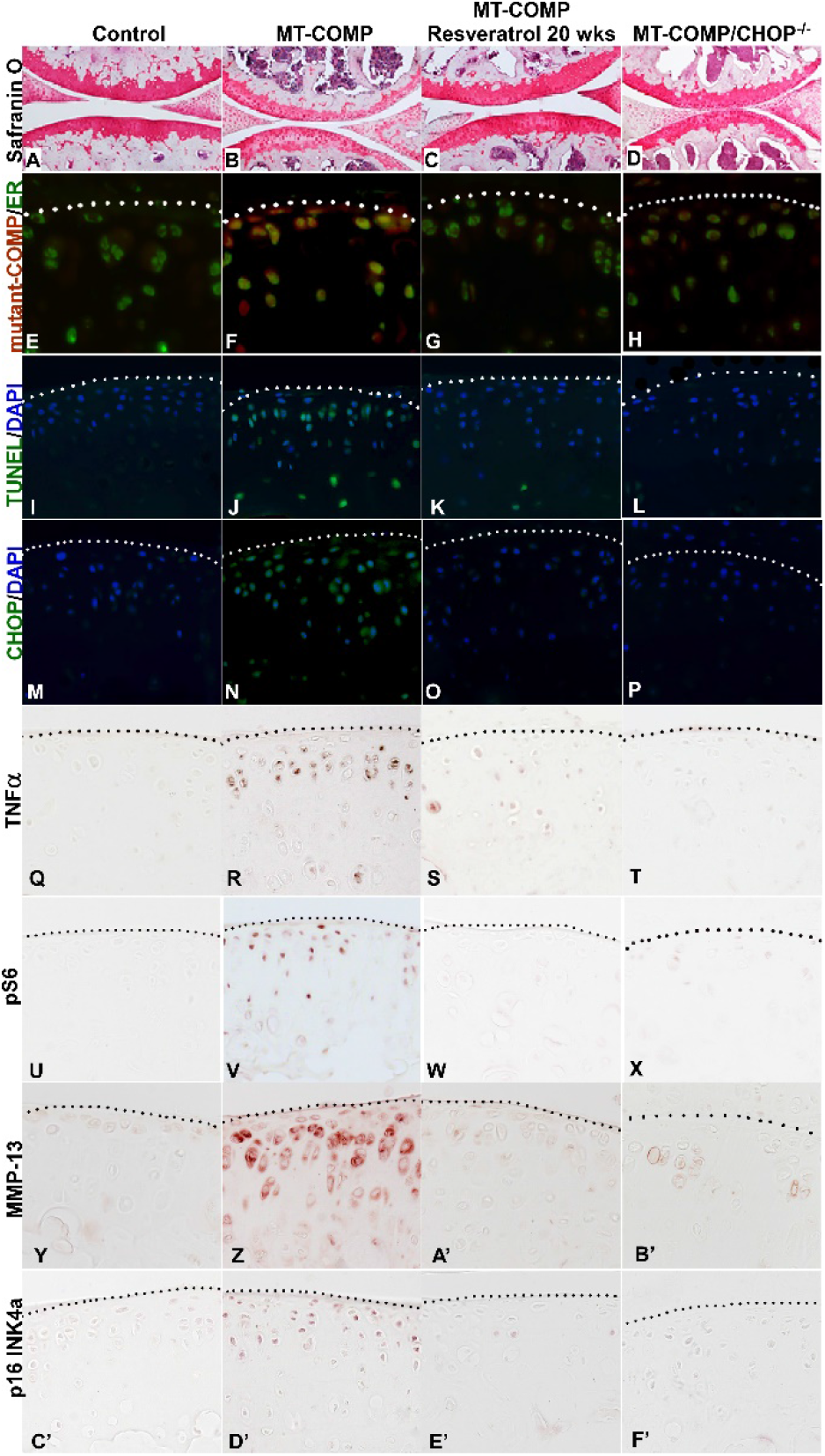
Articular cartilage of MT-COMP mice is preserved with resveratrol treatment or CHOP ablation. DOX was administered to all mice and resveratrol treatment to one group of MT-COMP mice (C,G,K,O,S,W,A’,E’) from birth to 20 wk. All articular cartilages were stained with safranin O (**A-D**) or immunostained using human COMP-specific antibodies (red, **E-H**) and protein disulfide isomerase (PDI) an ER marker (green, **E-H**) or TUNEL (green, **I-L**) and DAPI (blue nuclei, **I-L**) or CHOP (green, **M-P** ER stress marker) and DAPI (blue, nuclei **M-P**), TNFα (**Q-T**), pS6 (**U-X**), MMP13 (**Y-B’**) or p16 INK4a (**C’-F’**). Safranin O staining shows that MT-COMP articular cartilage has less proteoglycans (**B**) compared to control (**A**) and both resveratrol treated and ablation of CHOP normalizes proteoglycan content (**C, D**). TUNEL positive MT-COMP articular chondrocytes in (**J**) are more numerous than in the control (**I**), resveratrol treated (**K**), or MT-COMP/CHOP^-/-^ (**L**). TNFα inflammation is decreased in resveratrol treated (**S**), or MT-COMP/CHOP^-/-^ (**T**) articular chondrocytes compared to MT-COMP (**R**). mTORC1 signal activity (pS6) is detected in MT-COMP mice (**V**) compared to controls (**U**), resveratrol treated MT-COMP (**W**) or MT-COMP/CHOP^-/-^ (**X**) mice. Increased MMP-13 is present in MT-COMP (**Z**) articular cartilage compared to controls (**Y**), resveratrol treated (**A’**) or MT-COMP/CHOP^-/-^ (**B’**) mice. Senescent articular chondrocytes (p16 INK4a) were observed in MT-COMP mice (**D’**), whereas minimal p16 INK4a signal is seen in controls (**C’**), resveratrol treated (**E’**) or MT-COMP/CHOP^-/-^ (**F’**). Bar =

## Discussion

Using our MT-COMP mouse model of PSACH, we found that retention of mutant-COMP in the ER of articular chondrocytes induces and drives a CHOP-dependent ER stress pathologic loop involving multiple inflammatory processes. Persistent inflammation drives autophagy blockage, and ultimately, chondrocyte death. The presence of senescence and a degenerative environment likely adversely impacts surrounding tissue including synovium and subchondral bone. The results of this study show that MT-COMP mice undergo premature joint degeneration similar to the premature joint degeneration in PSACH providing a model system for studying nonsurgical joint sparing therapeutics.

Multiple cellular stresses were observed in MT-COMP articular chondrocytes including ER stress (CHOP), inflammation (TNFα), ECM degrative enzymes (MMP-13), block of autophagy (pS6) and senescence (p16 INK4a). Importantly, these stresses were not observed in MT-COMP mice in the absence of DOX (**Supplemental Fig. 1**). TNFα stimulates excessive mTORC1 signaling (observed by pS6 immunostaining) blocking autophagy and preventing mutant-COMP from being cleared from the ER through macroautophagy (referred to as autophagy). The autophagy blockade means that the ER cannot be cleared and the pathology is perpetuated without therapeutic intervention. Moreover, elevated mTORC1 signaling maintains protein synthesis and this counteracts the unfolded protein response repression of translation to assist with clearance of the ER.

Aging and/or cellular stress stimulates senescence [62, 76] and senescent articular chondrocytes were observed in the articular cartilage of MT-COMP mice at 16 wks. The multiple chronic stresses that occur in MT-COMP articular chondrocytes likely stimulate senescence similar to that observed in OA [61]. Based on our ER-stress mechanistic model, we expect that inflammation, ER stress, MMP expression, autophagy blockage, senescence and chondrocyte death, each will contribute to articular cartilage erosion [45, 47–49, 77–79]. Unique to the MT-COMP articular chondrocytes is the presence of well-known degradative enzyme, MMP-13, and senescence both of which are associated with OA in humans and animal models [61, 80]. These findings suggest that mutant-COMP joint degeneration shares some cellular pathology with OA.

The timing of mutant-COMP joint degeneration is distinct from the mutant-COMP growth plate pathology. In articular chondrocytes, mutant-COMP retention and inflammation start at 4 wks and ER stress, blockage of autophagy and chondrocyte death are seen between 8 and 12 wks (summarized in **Table 2**). In contrast, mutant-COMP retention in the growth plate is seen prenatally with inflammation starting post-nally at 2 wks and peaking between 3-4 wks, and chondrocyte death significantly increased by 4 wks [39, 47, 48]. These tissues serve very different functions with growth plate driving linear growth from birth to 10 wks and the articular cartilage needed to cushion mechanical forces that are not required in early life (birth to 3 wks) when ambulation is limited. Articular cartilages primarily absorb and distribute mechanical forces so these forces are not transmitted to the bone. In contrast, the growth plate is a niche for the maturation of growth plate chondrocytes that eventually generate copious amount of ECM that will be calcified and turned into bone. These functional differences may explain the novel mutant-COMP articular chondrocyte pathology of matrix degradation, senescence and premature joint degeneration in MT-COMP mice.

**Table 2:** Timing of articular chondrocyte pathology. DOX administered from birth to collection at 2, 4, 8, 12, 16, 20 wks. Mutant-COMP intracellular retention, proteoglycan loss, ER stress, TNF inflammation, autophagy block, senescent chondrocytes, degradation enzyme MMP-13 and chondrocyte death are present (X) and begin between 4-16 wks. * = some immunosignal.

| **Mutant-COMP pathology** | **2** | **4** | **8** | **12** | **16** | **20 wks.** |
| --- | --- | --- | --- | --- | --- | --- |
| Mutant-COMP intracellular retention |  | X | X | X | X | X |
| Proteoglycan loss (Safranin O) |  |  |  |  | X | X |
| ER stress (CHOP) |  |  |  | X | X | X |
| Chondrocyte death (TUNEL) |  |  | X | X | X | X |
| Inflammation (TNFα) | * | X | X | X | X | X |
| Autophagy block (pS6) |  |  | * | X | X | X |
| Enzymatic degradation (MMP-13) |  |  | * | X | X | X |
| Senescent chondrocytes (p16 INK4a) |  |  |  | * | X | X |

MT-COMP mice undergo premature joint degeneration far earlier than the background strain (C57BL\6) that develop joint degeneration beginning at 1 year of age. While there is limited information about the temporal development of joint degeneration in PSACH, natural history studies show that there is premature joint degeneration in PSACH starting in mid-to late teenage years. [81]. Based on this information, we posit that the MT-COMP mouse is a good model system for understanding the PSACH/mutant-COMP pathologies.

The essential role of CHOP in the mutant-COMP joint degenerative process is illustrated by the repression of articular chondrocyte stress in MT-COMP/CHOP^-/-^mice. Notably, the MT-COMP/CHOP^-/-^mice still express mutant-COMP but in the absence of CHOP the mutant-COMP pathology is diminished. Similarly, resveratrol treatment from birth to 20 wks dampens mutant-COMP pathologies of ER stress (CHOP), inflammation (TNFα), block of autophagy (pS6), senescence (p16 INK4a). This reduction of ER stress and inflammation interrupts the pathological loop between ER, oxidative stress and inflammation. Moreover, resveratrol restoration of autophagy allows clearance of mutant-COMP from the ER of articular chondrocytes. These proof-of-principle findings show that the articular cartilage of MT-COMP mice can be preserved if intervention occurs prior to a point of no return and opens new therapeutic avenues for PSACH.

## Materials and Methods

### Bigenic mice

The MT-COMP mice used in these and previously described experiments contain the pTRE-COMP (coding sequence of human COMP+FLAG tag driven by the tetracycline responsive element promoter) and pTET-On-Col II (rtTA coding sequence driven by a type II collagen promoter) [39, 47, 53]. DOX (500 ng/ml) was administered to mice at birth (through mother’s milk) to collection in their drinking water. This study complied with the Guide for the Care and Use of Laboratory Animals, eighth edition (ISBN-10, 0-309-15396-4) and was approved by the Animal Welfare Committee at the University of Texas Medical School at Houston and complies with NIH guidelines.

### Generation of CHOP null bigenic mice

CHOP null mice were procured from Jackson Laboratories and mated with MT-COMP bigenic mice to obtain a strain expressing MT-COMP in a CHOP null background (MT-COMP/CHOP^-/-^) and these mice were used in previous experiments [47, 53]. Genotypes of the CHOP null mice were verified using CHOP-specific primers [53].

### Immunohistochemistry

Hind limbs from male and female MT-COMP and C57BL\6 control mice were collected and articular cartilage analyzed as previously described [39, 47, 79]. Briefly, the limbs were fixed in: 95% ethanol followed by decalcification in immunocal (StatLab McKinney, TX) for 1 week and pepsin (1 mg/ml in 0.1N HCl) was used for antigen retrieval for immunostaining with human COMP (Thermofisher; MA1-20221, 1:100), CHOP (Santa Cruz; SC-575; 1:100), interleukin 1 (IL-1) (Abcam; ab7632, 1:200), tumor necrosis factor α (TNFα) (Abcam; ab6671, 1:200), PDI (Santa Cruz; SC-20132, 1:100), p16 INK4a - (Abcam; ab189034, 1:200), pS6 (Cell Signaling Technology 2215S rabbit polyclonal, 1:200), MMP-13 (Abcam: ab39012, 1:50) or 10% wt/vol formalin for terminal deoxynucleotidyl transferase–mediated deoxyuridine triphosphate-biotin nick end (TUNEL) labeling. Species specific biotinylated secondary antibodies were used for 1 hr at RT. Sections of the same thickness were then washed and incubated with streptavidin horseradish peroxidase (HRP) and DAB was used as chromogen. The sections were dehydrated and mounted with Thermofisher cytoseal 60 and then visualized under the BX51 inverted microscope (Olympus America, Center Valley, PA). Limbs were fixed in 10% wt/vol formalin for terminal deoxynucleotidyl transferase–mediated deoxyuridine triphosphate-biotin nick end labeling (TUNEL) staining. For proteoglycan stains, samples were deparaffinized and hydrated in distilled water and stained with safranin-O (Spectrum Chemical, New Brunswick, NJ, 477-73-6) according to manufacturer’s protocol. Immunostaining was performed on 10 animals in each group.

### Gait analysis

Gait was analyzed using a DigiGait treadmill system (Mouse Specifics Inc., Boston, MA) following the protocol described for collagen-induced OA [39]. Previously, it has been shown that running during gait analysis was necessary to uncover subtle changes in ambulatory function that are associated with OA [73]. Briefly, video camera images of mice running through a transparent belt were captured and analyzed. At least 3 sec of downhill 15° running continuous strides were used to calculate and measure gait parameters with DigiGait software. Mice were run for 30 cm/sec and Gait analysis was performed in the animal behavioral testing room and assessments were performed by software so blinding was not necessary.

DigiGait recommends 3 mice per group to measure ethanol induced ataxia or running speed (https://mousespecifics.com/digitgait-protocols/-lastaccessed5-22-21). Based on this information, 5 mice/group were analyzed because OA gait dysfunction is subtler to detect compared to ethanol induced ataxia. ANOVA analysis was performed followed by pairwise comparisons with Kruskal-Wallis and post hoc Dunn’s analysis. All mice were moved to the behavior room at 8:00 am and DigiGait analysis occurred between 9:00 am - 12:30 pm.

### OA scoring

OA scoring was performed on 10 different sections (from individual mice) divided into quadrants (medial femoral condyle, medial tibial plateau, lateral femoral condyle, lateral tibial plateau). Scores ranged from 0 for normal cartilage to 6 for lesions reaching the calcified cartilage for >75% of the quadrant width and are presented as a total score and each score listed individually [82]. While OARSI scoring covers a wide range of OA pathology, in this study, OA scoring was modified to optimize evaluation of early OA pathology. Four areas, synovium, bone/cartilage, tibial and femoral articular cartilage, were scored from 0 to 3 on safranin-O stained sections. A score of 0 indicated normal or no damage, 1 = mild damage, 2 = moderate damage and 3 = denotes severe damage. Synovitis, bone/cartilage damage and proteoglycan of tibia and femur were scored individually and all scores summed for a with a maximal score of 12. <u>Synovitis</u> was defined as: mild - increase in thickness of synovial lining and increase in stromal area, moderate - increase in stromal density or severe - synovial lining is thickened with further increase of stromal cellular density. <u>Bone/cartilage damage</u> defined as: normal - surface was smooth, mild −minor erosion of the surface, moderate −presence of remodeling with minor erosion, or severe −major erosion. <u>Proteoglycans</u> of the articular cartilage of the <u>tibia</u> and <u>femur</u> was classified as: normal −if staining was even through to the subchondral bone, mild -when staining was thinned, moderate - with thinning of proteoglycan stained layer and absence of staining in some areas or severe - widespread loss of proteoglycan staining. Ten mice per experimental group were used for each time point providing 80-90% power to detect a minimal difference of 2 or 3 units. All scoring was performed blindly. Section depth, thickness, fixation, decalcification conditions were all identical for all limbs analyzed. Kruskal-Wallis test was used to evaluate distribution of OA scores across at 6 experiment groups with Post-hoc Dunn’s test comparing MT-COMP to controls.

#### MMPSense

Mice were treated with depilatory cream to remove hair prior to imaging. MMP 680 reagent was injected in tail vein and animals and imaged 24 hrs. later per manufacturer’s instructions with an uninjected control (UIC) and imaged on an IVIS Spectrum *In Vivo* Imaging System (PerkinElmer; Waltham, MA) (https://www.perkinelmer.com/lab-solutions/resources/docs/APP_Protocol_MMPSense%20680.pdf last accessed 2-22-21). MMP 680 is an optically silent substrate which fluoresces when cleaved by MMP−2, −3, −9, and −13. MMPSense signal was assessed blindly by positioning a circle of a standard size around the knee (in all samples) and radiance efficiency was generated from IVIS software. Six mice were included per group and had the power to detect a difference of 30% or greater. Mann-Whitney U test was used to evaluate MMP activity in control and MT-COMP mice.

## Acknowledgements

This work was supported by the National Institute of Arthritis and Musculoskeletal and Skin Diseases of the National Institutes of Health (NIAMS) [Award Number 5R01AR057117-10], the Leah Lewis Family Foundation and The Rolanette and Berdon Lawrence Bone Disease Program of Texas. The content is solely the responsibility of the authors and does not necessarily represent the official views of the National Institutes of Health. We thank Dr. Joseph L. Alcorn for manuscript editing assistance and helpful discussions.

## Author contributions

Conceptualization, K.L.P and J.T.H.; Methodology, A.C.V and D.P.; Formal Analysis, K.L.P; Investigation, A.C.V, M.G.H, F.C., D.P. and F.C.; Resources, J.T.H.; Data Curation, F.C., M.G.H, F.C. and D.P.; Writing – Original Draft Preparation, K.L.P.; Writing – Review & Editing, K.L.P and J.T.H.; Supervision, K.L.P and J.T.H.; Project Administration, K.L.P and J.T.H.; Funding Acquisition, K.L.P and J.T.H.

## Conflicts of Interest

The authors declare no conflict of interest.

---

**Supplemental Figure 1:**
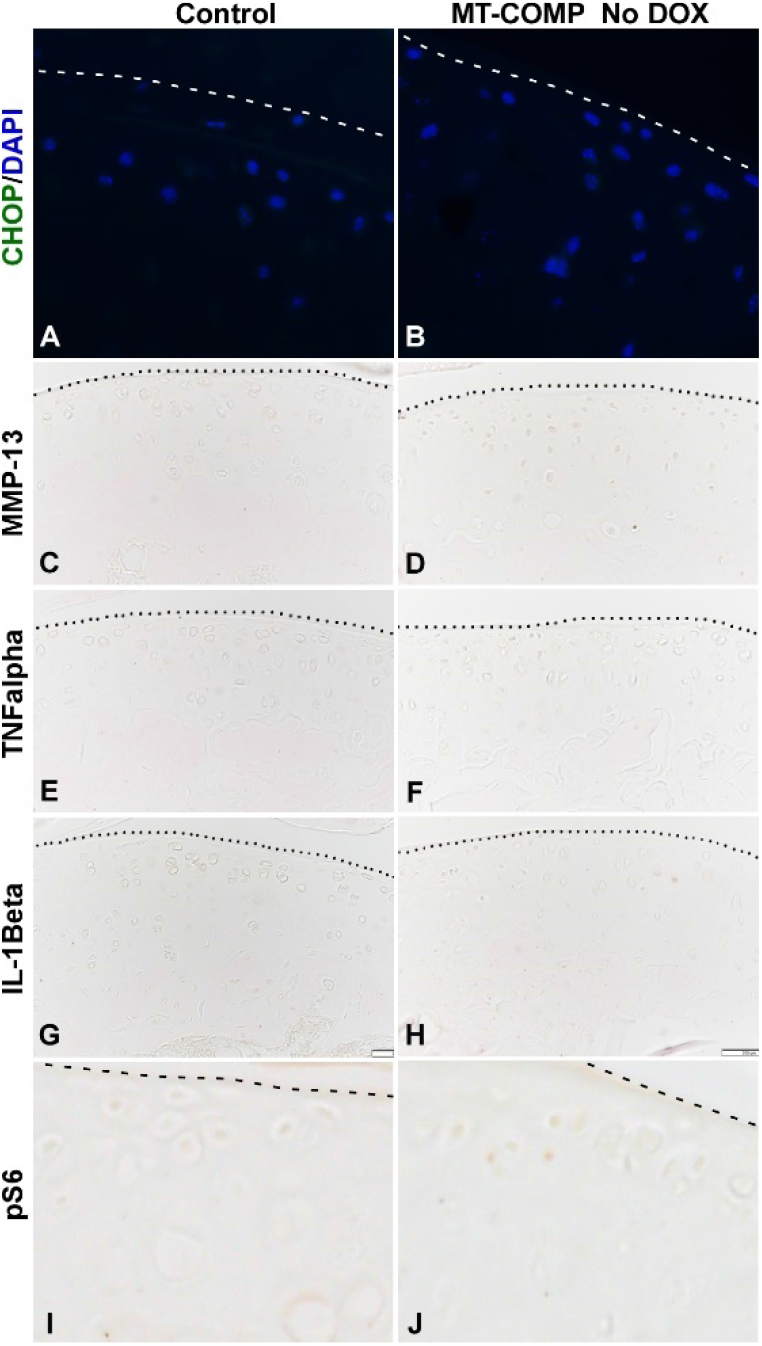
Absence of mutant-COMP expression correlates with no articular chondrocyte stress in MT-COMP mice. Articular cartilage from control and MT-COMP mice with no DOX were immunostained for CHOP (green, **A-B** ER) and DAPI (blue nuclei, **A-B**) or MMP13 (**C-D**) (green **I-L**) or TNFα (**E-F**) of IL-1β (**G-H**) or pS6 (**I-J**). ER stress (CHOP), degradative enzyme (MMP13), inflammation (TNFα or IL-1β) and block of autophagy (pS6) were not detected in either MT-COMP or control articular chondrocytes in the absence of DOX that induces mutant-COMP expression.

## Notes

### Competing Interest Statement

The authors have declared no competing interest.

## References

1. DiCesare, P. E.; Morgelin, M.; Mann, K.; Paulsson, M., Cartilage oligomeric matrix protein and thrombospondin 1. Purification from articular cartilage, electron microscopic structure, and chondrocyte binding. Eur J Biochem 1994, 223, (3), 927–37.

2. Hecht, J. T.; Deere, M.; Putnam, E.; Cole, W.; Vertel, B.; Chen, H.; Lawler, J., Characterization of cartilage oligomeric matrix protein (COMP) in human normal and pseudoachondroplasia musculoskeletal tissues. Matrix Biol 1998, 17, (4), 269–78.

3. Hedbom, E.; Antonsson, P.; Hjerpe, A.; Aeschlimann, D.; Paulsson, M.; Rosa-Pimentel, E.; Sommarin, Y.; Wendel, M.; Oldberg, A.; Heinegard, D., Cartilage matrix proteins. An acidic oligomeric protein (COMP) detected only in cartilage. J Biol Chem 1992, 267, (9), 6132–6.

4. Urban, J. P.; Maroudas, A.; Bayliss, M. T.; Dillon, J., Swelling pressures of proteoglycans at the concentrations found in cartilaginous tissues. Biorheology 1979, 16, (6), 447–64.

5. Kempson, G. E.; Freeman, M. A.; Swanson, S. A., Tensile properties of articular cartilage. Nature 1968, 220, (172), 1127–8.

6. Schmidt, M. B.; Mow, V. C.; Chun, L. E.; Eyre, D. R., Effects of proteoglycan extraction on the tensile behavior of articular cartilage. Journal of Orthopaedic Research 1990, 8, (3), 353–63.

7. Adams, J. C.; Tucker, R. P.; Lawler, J., The thrombospondin gene family. R.G. Landes Co.: Austin, Tex., U.S.A., 1995; p 191.

8. Holden, P.; Meadows, R. S.; Chapman, K. L.; Grant, M. E.; Kadler, K. E.; Briggs, M. D., Cartilage oligomeric matrix protein interacts with type IX collagen, and disruptions to these interactions identify a pathogenetic mechanism in a bone dysplasia family. J Biol Chem 2001, 276, (8), 6046–55.

9. Thur, J.; Rosenberg, K.; Nitsche, D. P.; Pihlajamaa, T.; Ala-Kokko, L.; Heinegard, D.; Paulsson, M.; Maurer, P., Mutations in cartilage oligomeric matrix protein causing pseudoachondroplasia and multiple epiphyseal dysplasia affect binding of calcium and collagen I, II, and IX. J Biol Chem 2001, 276, (9), 6083–92.

10. Chen, F. H.; Herndon, M. E.; Patel, N.; Hecht, J. T.; Tuan, R. S.; Lawler, J., Interaction of cartilage oligomeric matrix protein/thrombospondin 5 with aggrecan. J Biol Chem 2007, 282, (34), 24591–8.

11. Mann, H. H.; Ozbek, S.; Engel, J.; Paulsson, M.; Wagener, R., Interactions between the cartilage oligomeric matrix protein and matrilins. Implications for matrix assembly and the pathogenesis of chondrodysplasias. J Biol Chem 2004, 279, (24), 25294–8.

12. Di Cesare, P. E.; Chen, F. S.; Moergelin, M.; Carlson, C. S.; Leslie, M. P.; Perris, R.; Fang, C., Matrix-matrix interaction of cartilage oligomeric matrix protein and fibronectin. Matrix Biol 2002, 21, (5), 461–70.

13. Smith, R. K.; Gerard, M.; Dowling, B.; Dart, A. J.; Birch, H. L.; Goodship, A. E., Correlation of cartilage oligomeric matrix protein (COMP) levels in equine tendon with mechanical properties: a proposed role for COMP in determining function-specific mechanical characteristics of locomotor tendons. Equine Vet J Suppl 2002, (34), 241–4.

14. Kipnes, J.; Carlberg, A. L.; Loredo, G. A.; Lawler, J.; Tuan, R. S.; Hall, D. J., Effect of cartilage oligomeric matrix protein on mesenchymal chondrogenesis in vitro. Osteoarthritis Cartilage 2003, 11, (6), 442–54.

15. Xu, K.; Zhang, Y.; Ilalov, K.; Carlson, C. S.; Feng, J. Q.; Di Cesare, P. E.; Liu, C. J., Cartilage oligomeric matrix protein associates with granulin-epithelin precursor (GEP) and potentiates GEP-stimulated chondrocyte proliferation. J Biol Chem 2007, 282, (15), 11347–55.

16. Briggs, M. D.; Chapman, K. L., Pseudoachondroplasia and multiple epiphyseal dysplasia: mutation review, molecular interactions, and genotype to phenotype correlations. Hum Mutat 2002, 19, (5), 465–78.

17. Briggs, M. D.; Hoffman, S. M.; King, L. M.; Olsen, A. S.; Mohrenweiser, H.; Leroy, J. G.; Mortier, G. R.; Rimoin, D. L.; Lachman, R. S.; Gaines, E. S., Pseudoachondroplasia and multiple epiphyseal dysplasia due to mutations in the cartilage oligomeric matrix protein gene. Nature Genetics 1995, 10, (3), 330–6.

18. Briggs, M. D.; Mortier, G. R.; Cole, W. G.; King, L. M.; Golik, S. S.; Bonaventure, J.; Nuytinck, L.; De Paepe, A.; Leroy, J. G.; Biesecker, L.; Lipson, M.; Wilcox, W. R.; Lachman, R. S.; Rimoin, D. L.; Knowlton, R. G.; Cohn, D. H., Diverse mutations in the gene for cartilage oligomeric matrix protein in the pseudoachondroplasia-multiple epiphyseal dysplasia disease spectrum. Am J Hum Genet 1998, 62, (2), 311–9.

19. Chen, H.; Deere, M.; Hecht, J. T.; Lawler, J., Cartilage oligomeric matrix protein is a calcium-binding protein, and a mutation in its type 3 repeats causes conformational changes. J Biol Chem 2000, 275, (34), 26538–44.

20. Chen, T. L.; Stevens, J. W.; Cole, W. G.; Hecht, J. T.; Vertel, B. M., Cell-type specific trafficking of expressed mutant COMP in a cell culture model for PSaCh. Matrix Biol 2004, 23, (7), 433–44.

21. Cooper, R. R.; Ponseti, I. V.; Maynard, J. A., Pseudoachondroplastic dwarfism. A rough-surfaced endoplasmic reticulum storage disorder. Journal of Bone & Joint Surgery - American Volume 1973, 55, (3), 475–84.

22. Delot, E.; Brodie, S. G.; King, L. M.; Wilcox, W. R.; Cohn, D. H., Physiological and pathological secretion of cartilage oligomeric matrix protein by cells in culture. J Biol Chem 1998, 273, (41), 26692–7.

23. DiCesare, P. E.; Morgelin, M.; Carlson, C. S.; Pasumarti, S.; Paulsson, M., Cartilage oligomeric matrix protein: isolation and characterization from human articular cartilage. J Orthop Res 1995, 13, (3), 422–8.

24. Dinser, R.; Zaucke, F.; Kreppel, F.; Hultenby, K.; Kochanek, S.; Paulsson, M.; Maurer, P., Pseudoachondroplasia is caused through both intra- and extracellular pathogenic pathways. J Clin Invest 2002, 110, (4), 505–13.

25. Duke, J.; Montufar-Solis, D.; Underwood, S.; Lalani, Z.; Hecht, J. T., Apoptosis staining in cultured pseudoachondroplasia chondrocytes. Apoptosis 2003, 8, (2), 191–7.

26. Ikegawa, S.; Ohashi, H.; Nishimura, G.; Kim, K. C.; Sannohe, A.; Kimizuka, M.; Fukushima, Y.; Nagai, T.; Nakamura, Y., Novel and recurrent COMP (cartilage oligomeric matrix protein) mutations in pseudoachondroplasia and multiple epiphyseal dysplasia. Hum Genet 1998, 103, (6), 633–8.

27. Kleerekoper, Q.; Hecht, J. T.; Putkey, J. A., Disease-causing mutations in cartilage oligomeric matrix protein cause an unstructured Ca2+ binding domain. J Biol Chem 2002, 277, (12), 10581–9.

28. Maddox, B. K.; Mokashi, A.; Keene, D. R.; Bachinger, H. P., A cartilage oligomeric matrix protein mutation associated with pseudoachondroplasia changes the structural and functional properties of the type 3 domain. J Biol Chem 2000, 275, (15), 11412–7.

29. McKeand, J.; Rotta, J.; Hecht, J. T., Natural history study of pseudoachondroplasia. Am J Med Genet 1996, 63, (2), 406–10.

30. Merritt, T. M.; Alcorn, J. L.; Haynes, R.; Hecht, J. T., Expression of mutant cartilage oligomeric matrix protein in human chondrocytes induces the pseudoachondroplasia phenotype. J Orthop Res 2006, 24, (4), 700–7.

31. Merritt, T. M.; Bick, R.; Poindexter, B. J.; Alcorn, J. L.; Hecht, J. T., Unique Matrix Structure in the Rough Endoplasmic Reticulum Cisternae of Pseudoachondroplasia Chondrocytes. The American Journal of Pathology 2007, 170, (1), 293–300.

32. Unger, S.; Hecht, J. T., Pseudoachondroplasia and multiple epiphyseal dysplasia: New etiologic developments. American Journal of Medical Genetics 2001, 106, (4), 244.

33. Hecht, J. T.; Montufar-Solis, D.; Decker, G.; Lawler, J.; Daniels, K.; Duke, P. J., Retention of cartilage oligomeric matrix protein (COMP) and cell death in redifferentiated pseudoachondroplasia chondrocytes. Matrix Biology 1998, 17, (8-9), 625–633.

34. Hecht, J. T.; Hayes, E.; Haynes, R.; Cole, W. G., COMP mutations, chondrocyte function and cartilage matrix. Matrix Biol 2005, 23, (8), 525–33.

35. Hecht, J. T.; Hayes, E.; Snuggs, M.; Decker, G.; Montufar-Solis, D.; Doege, K.; Mwalle, F.; Poole, R.; Stevens, J.; Duke, P. J., Calreticulin, PDI, Grp94 and BiP chaperone proteins are associated with retained COMP in pseudoachondroplasia chondrocytes. Matrix Biol 2001, 20, (4), 251–62.

36. Hecht, J. T.; Nelson, L. D.; Crowder, E.; Wang, Y.; Elder, F. F.; Harrison, W. R.; Francomano, C. A.; Prange, C. K.; Lennon, G. G.; Deere, M.; Lawler, J., Mutations in exon 17B of cartilage oligomeric matrix protein (COMP) cause pseudoachondroplasia. Nat Genet 1995, 10, (3), 325–9.

37. Posey, K. L.; Hayes, E.; Haynes, R.; Hecht, J. T., Role of TSP-5/COMP in pseudoachondroplasia. Int J Biochem Cell Biol 2004, 36, (6), 1005–12.

38. Bonafe, L.; Cormier-Daire, V.; Hall, C.; Lachman, R.; Mortier, G.; Mundlos, S.; Nishimura, G.; Sangiorgi, L.; Savarirayan, R.; Sillence, D.; Spranger, J.; Superti-Furga, A.; Warman, M.; Unger, S., Nosology and classification of genetic skeletal disorders: 2015 revision. Am J Med Genet A 2015, 167A, (12), 2869–92.

39. Posey, K. L.; Veerisetty, A. C.; Liu, P.; Wang, H. R.; Poindexter, B. J.; Bick, R.; Alcorn, J. L.; Hecht, J. T., An Inducible Cartilage Oligomeric Matrix Protein Mouse Model Recapitulates Human Pseudoachondroplasia Phenotype. The American Journal of Pathology 2009, 175, (4), 1555–1563.

40. Maynard, J. A.; Cooper, R. R.; Ponseti, I. V., A unique rough surfaced endoplasmic reticulum inclusion in pseudoachondroplasia. Lab Invest 1972, 26, (1), 40–4.

41. Hall, J. G., Pseudoachondroplasia. Birth Defects Orig Artic Ser 1975, 11, (6), 187–202.

42. Posey, K. L.; Hecht, J. T., The role of cartilage oligomeric matrix protein (COMP) in skeletal disease. Curr Drug Targets 2008, 9, (10), 869–77.

43. Cooper, R. R.; Ponseti, I. V.; Maynard, J. A., Pseudoachondroplasia dwarfism. A rough-surfaced endoplasmic reticulum disorder. J Bone Joint Surg Am 1973, 55A, 475–484.

44. Kvansakul, M.; Adams, J. C.; Hohenester, E., Structure of a thrombospondin C-terminal fragment reveals a novel calcium core in the type 3 repeats. Embo J 2004, 23, (6), 1223–33.

45. Posey, K. L.; Hecht, J. T., Novel therapeutic interventions for pseudoachondroplasia. Bone 2017.

46. Coustry, F.; Posey, K. L.; Liu, P.; Alcorn, J. L.; Hecht, J. T., D469del-COMP Retention in Chondrocytes Stimulates Caspase-Independent Necroptosis. The American Journal of Pathology 2012, 180, (2), 738–748.

47. Posey, K. L.; Coustry, F.; Veerisetty, A. C.; Liu, P.; Alcorn, J. L.; Hecht, J. T., Chop (Ddit3) is essential for D469del-COMP retention and cell death in chondrocytes in an inducible transgenic mouse model of pseudoachondroplasia. Am J Pathol 2012, 180, (2), 727–37.

48. Posey, K. L.; Coustry, F.; Veerisetty, A. C.; Liu, P.; Alcorn, J. L.; Hecht, J. T., Chondrocyte-specific pathology during skeletal growth and therapeutics in a murine model of pseudoachondroplasia. J Bone Miner Res 2014, 29, (5), 1258–68.

49. Posey, K. L.; Coustry, F.; Veerisetty, A. C.; Hossain, M. G.; Gambello, M. J.; Hecht, J. T., Novel mTORC1 Mechanism Suggests Therapeutic Targets for COMPopathies. Am J Pathol 2019, 189, (1), 132–146.

50. Decker, R. S.; Koyama, E.; Pacifici, M., Articular Cartilage: Structural and Developmental Intricacies and Questions. Curr Osteoporos Rep 2015, 13, (6), 407–14.

51. Mangiavini, L.; Merceron, C.; Schipani, E., Analysis of Mouse Growth Plate Development. Curr Protoc Mouse Biol 2016, 6, (1), 67–130.

52. Maroteaux, P.; Lamy, M., [Pseudo-achondroplastic forms of spondylo-epiphyseal dysplasias.]. Presse Med 1959, 67, (10), 383–6.

53. Marciniak, S. J.; Yun, C. Y.; Oyadomari, S.; Novoa, I.; Zhang, Y.; Jungreis, R.; Nagata, K.; Harding, H. P.; Ron, D., CHOP induces death by promoting protein synthesis and oxidation in the stressed endoplasmic reticulum. Genes Dev 2004, 18, (24), 3066–77.

54. Burrage, P. S.; Mix, K. S.; Brinckerhoff, C. E., Matrix metalloproteinases: role in arthritis. Front Biosci 2006, 11, 529–43.

55. Goldring, M. B.; Otero, M., Inflammation in osteoarthritis. Curr Opin Rheumatol 2011, 23, (5), 471–8.

56. Goldring, M. B., Articular cartilage degradation in osteoarthritis. HSS J 2012, 8, (1), 7–9.

57. Jeon, H.; Im, G. I., Autophagy in osteoarthritis. Connect Tissue Res 2017, 58, (6), 497–508.

58. Rabanal-Ruiz, Y.; Otten, E. G.; Korolchuk, V. I., mTORC1 as the main gateway to autophagy. Essays Biochem 2017, 61, (6), 565–584.

59. Rim, Y. A.; Nam, Y.; Ju, J. H., The Role of Chondrocyte Hypertrophy and Senescence in Osteoarthritis Initiation and Progression. Int J Mol Sci 2020, 21, (7).

60. Bolduc, J. A.; Collins, J. A.; Loeser, R. F., Reactive oxygen species, aging and articular cartilage homeostasis. Free Radic Biol Med 2019, 132, 73–82.

61. Coryell, P. R.; Diekman, B. O.; Loeser, R. F., Mechanisms and therapeutic implications of cellular senescence in osteoarthritis. Nat Rev Rheumatol 2021, 17, (1), 47–57.

62. Kumari, R.; Jat, P., Mechanisms of Cellular Senescence: Cell Cycle Arrest and Senescence Associated Secretory Phenotype. Front Cell Dev Biol 2021, 9, 645593.

63. Loeser, R. F., Aging and osteoarthritis: the role of chondrocyte senescence and aging changes in the cartilage matrix. Osteoarthritis Cartilage 2009, 17, (8), 971–9.

64. Calvo, E.; Palacios, I.; Delgado, E.; Sanchez-Pernaute, O.; Largo, R.; Egido, J.; Herrero-Beaumont, G., Histopathological correlation of cartilage swelling detected by magnetic resonance imaging in early experimental osteoarthritis. Osteoarthritis Cartilage 2004, 12, (11), 878–86.

65. Madry, H.; Luyten, F. P.; Facchini, A., Biological aspects of early osteoarthritis. Knee Surg Sports Traumatol Arthrosc 2012, 20, (3), 407–22.

66. Ruan, M. Z.; Erez, A.; Guse, K.; Dawson, B.; Bertin, T.; Chen, Y.; Jiang, M. M.; Yustein, J.; Gannon, F.; Lee, B. H., Proteoglycan 4 expression protects against the development of osteoarthritis. Sci Transl Med 2013, 5, (176), 176ra34.

67. Wei, L.; Fleming, B. C.; Sun, X.; Teeple, E.; Wu, W.; Jay, G. D.; Elsaid, K. A.; Luo, J.; Machan, J. T.; Chen, Q., Comparison of differential biomarkers of osteoarthritis with and without posttraumatic injury in the Hartley guinea pig model. J Orthop Res 2010, 28, (7), 900–6.

68. Yin, W.; Park, J. I.; Loeser, R. F., Oxidative stress inhibits insulin-like growth factor-I induction of chondrocyte proteoglycan synthesis through differential regulation of phosphatidylinositol 3-Kinase-Akt and MEK-ERK MAPK signaling pathways. J Biol Chem 2009, 284, (46), 31972–81.

69. Deuis, J. R.; Dvorakova, L. S.; Vetter, I., Methods Used to Evaluate Pain Behaviors in Rodents. Front Mol Neurosci 2017, 10, 284.

70. Sheahan, T. D.; Copits, B. A.; Golden, J. P.; Gereau, R. W. t., Voluntary Exercise Training: Analysis of Mice in Uninjured, Inflammatory, and Nerve-Injured Pain States. PLoS One 2015, 10, (7), e0133191.

71. Cobos, E. J.; Portillo-Salido, E., “Bedside-to-Bench” Behavioral Outcomes in Animal Models of Pain: Beyond the Evaluation of Reflexes. Curr Neuropharmacol 2013, 11, (6), 560–91.

72. Lakes, E. H.; Allen, K. D., Gait analysis methods for rodent models of arthritic disorders: reviews and recommendations. Osteoarthritis Cartilage 2016, 24, (11), 1837–1849.

73. Jacobs, B. Y.; Kloefkorn, H. E.; Allen, K. D., Gait analysis methods for rodent models of osteoarthritis. Curr Pain Headache Rep 2014, 18, (10), 456.

74. Kwok, J.; Onuma, H.; Olmer, M.; Lotz, M. K.; Grogan, S. P.; D’Lima, D. D., Histopathological analyses of murine menisci: implications for joint aging and osteoarthritis. Osteoarthritis Cartilage 2016, 24, (4), 709–18.

75. Hecht, J. T.; Coustry, F.; Veerisetty, A. C.; Hossain, M. G.; Posey, K. L., Resveratrol Reduces COMPopathy in Mice Through Activation of Autophagy. JBMR Plus 2021, 5, (3), e10456.

76. van Deursen, J. M., The role of senescent cells in ageing. Nature 2014, 509, (7501), 439–46.

77. Posey, K. L.; Alcorn, J. L.; Hecht, J. T., Pseudoachondroplasia/COMP - translating from the bench to the bedside. Matrix Biol 2014, 37, 167–73.

78. Posey, K. L.; Coustry, F.; Hecht, J. T., Cartilage oligomeric matrix protein: COMPopathies and beyond. Matrix Biol 2018, 71-72, 161–173.

79. Posey, K. L.; Coustry, F.; Veerisetty, A. C.; Hossain, M.; Alcorn, J. L.; Hecht, J. T., Antioxidant and anti-inflammatory agents mitigate pathology in a mouse model of pseudoachondroplasia. Hum Mol Genet 2015, 24, (14), 3918–28.

80. Mehana, E. E.; Khafaga, A. F.; El-Blehi, S. S., The role of matrix metalloproteinases in osteoarthritis pathogenesis: An updated review. Life Sci 2019, 234, 116786.

81. Wilhelmi, G.; Ochsner, K.; Witzemann, E., [Etiopathogenetic significance of bone changes in spontaneous gonarthritis in the mouse]. Z Rheumatol 1986, 45, (1), 7–15.

82. Glasson, S. S.; Chambers, M. G.; Van Den Berg, W. B.; Little, C. B., The OARSI histopathology initiative - recommendations for histological assessments of osteoarthritis in the mouse. Osteoarthritis and Cartilage 2010, 18 Suppl 3, S17–23.

